# Causality in the natural selection of the population ecological life histories of birds

**DOI:** 10.1101/2024.03.23.586398

**Authors:** Lars Witting

## Abstract

Contingent life history theory explains evolution backwards by analysing the fitness consequences of tradeoffs and constraints in the evolved species of today, bypassing the essential challenge of predicting evolution forwardly by the cause and effect of natural selection. I do the latter to decompose the population ecological life histories of 11,187 species of birds. This shows how the selection of mass accounts for inter-specific variation, with 76% of the within order variance, and 72% of the between order differences, in the body mass, demography, and population ecological traits being reconciled by the response of population dynamic feedback selection to variation in net energy, mortality, and intra-specific interactive competition.

## 1 Introduction

With body masses that span four orders of magnitude, a pace of life that ranges from fast hummingbirds to slow breeding albatrosses, an average annual clutch size of less than one to more than twenty, and lifespans that last from a few to more than hundred years, birds have a tremendous amount of life history variation. Evolutionary theory aim to explain the variation, but contingent life history theory may confound trait correlations for natural selection causalities (Reeve and Sherman 2001; Witting 2008).

To get evolutionary causality right, it is useful to consider how natural selection consolidates itself in a hierarchical structure over time, allowing for the evolution of an increasing number of traits. This begins at the origin of self-replicating entities, where a relatively small set of traits can be selected independently of the derived traits that evolve later. A next natural selection level may emerge from the causal effects that the evolution of these independent traits has on the natural selection of other traits and/or trait components. Some of these trait-dependent components may be new traits that emerge from a selection imposed by the independent traits, while others may be fractions or components of the independent traits that were selected initially, being adjusted secondarily by their own evolutionary unfolding. These trait-dependent components may also be selected from ecologically induced changes in traits like mortality and resource consumption, even when the ecological changes are not themselves directly part of natural selection. This hierarchical linking of naturally selected traits continues during the course of evolution, generating higher level causalities that entangle the complete life history in a complex web of natural selection interactions.

Disentangling and decomposing this hierarchical structure of natural selection is one of the main goals of evolutionary studies. This work has taken many routes, with classical life history theory being the most widely applied (e.g. Lack 1947; Charlesworth 1980, 1994; Harvey and Pagel 1991; Roff 1992, 2002; Stearns 1992, Charnov 1993; Sibly et al. 2012). This theory developed during the 20th century focus on contingent evolution by historical natural selection (Witting 2008), following the principle that the evolution of biological organisation cannot be predicted forward even with full knowledge of antecedent conditions, but can only be understood backwards after time’s actual unfolding (Gould 2002; also Mayr 1988; Salthe 1989).

Based on the view that forward predictions are basically impossible, classical life history theory is primarily studying the natural selection hierarchy top-down backwards by analysing the fitness effects of trade-offs and constraints in the evolved species of today. This method uses mathematical life history models that explain the natural selection of dependent traits from traits and trade-offs that are treated as independent traits. And being developed from the population genetic synthesis (Fisher 1930; Wright 1931; Haldane 1932), the majority of classical life history theory operates by a frequency-independent selection where constant relative fitnesses are assigned to underlying genes.

Both the success and limitations of classical life history theory are tightly connected to this contingency. By measuring and modelling the fitness relations in evolved species, the approach is able to examine the evolutionary interactions among the different life history traits in real species. Yet, by developing and confirming evolutionary hypotheses from the evolved endpoints of current species, the approach is unsuited for a bottom-up forward analysis of the unfolding of the natural selection pressure that caused the evolution of the studied species in the first place. This limitation is reflected in mathematical models that do not address whether the assumed independent traits are indeed maintained by a natural selection that is more fundamental than the selection that explains the dependent traits. By not documenting such a bottom-up stability of natural selection, there is no guarantee that the real natural selection causality is not the opposite of the proposed, or a completely different, where the observed fitness interactions between traits are selected from primary variation in other factors (Witting 1997, 2008).

The theory of Malthusian relativity was constructed to document the bottom-up unfolding of natural selection by applying the alternative principle that variation in both the independent and dependent traits needs to be explained by the same natural selection model (Witting 1997, 2008). As the independent traits of the dependent traits may depend on the natural selection of a deeper layer of independent traits, this stronger theoretical principle cascades into a paradigm of inevitable evolution by deterministic natural selection. Given an abiotic environment suitable for life, inevitable evolution is the subset of biological evolution that follows from the natural selection that unfolds mechanistically from the origin of replicating molecules.

Being constructed as a bottom-up forward process, Malthusian relativity is well suited for studies on the buildup of natural selection causalities/pressures. There is however an upper limit to the number of natural selection interactions that can feasibly be handled by a single model, and the method is difficult to apply for analyses of complex fitness interactions at the evolved end-points of current species. But this limitation reflects the strength of a method that was constructed to use no evolved traits to explain as much as possible of the observed life history variation at all evolutionary scales.

While initially used to predict the evolution of body mass allometries in animals (Witting 1995), Malthusian relativity soon developed into a general theory of evolution based on the population dynamic feedback selection of the intra-specific density dependent interactive competition (Witting 1997). This selection follows from the origin of replicating molecules, where the selection of metabolism, body mass, and inter-specific allometries generates a gradual unfolding of the feedback; a unfolding that selects major lifeforms from replicating molecules, over prokaryote-like self-replicating cells, and larger unicells, to multicellular sexually reproducing animals (Witting 2002, 2017a,b).

In a recent study I integrated the allometric component of Malthusian relativity with almost 40,000 estimates of life history and ecological traits in birds, estimating population ecological life history models for 11,187 species of birds (Witting 2024). With the current paper, I use the trait variation of these models to show how the majority of the inter-specific variation in the body masses, life histories, and ecological traits of birds are reconciled by the inter-specific variation in a few traits at the core of population dynamic feed-back selection.

### 1.1 Bottom-up unfolding natural selection

In my analysis I decompose the inter-specific life history variation from a few primary drivers of population dynamic feedback selection, and this section identifies these main drivers.

Although most life history traits show a strong interspecific correlation with mass, body mass is not the primary driver of natural selection because its selection depends on other traits. Body mass is part of the quality-quantity trade-off (Smith and Fretwell 1974; Stearns 1992)—where a given amount of energy can produce a few large or many small offspring—and this selects for a continued decline in mass, when other things are equal.

The selection of mass in multicellular organisms is therefore dependent on an interactive competition where mass is selected as a competitive trait that is used by the larger than average individuals to monopolise resources. This occurs by a density-frequency-dependent selection, where the level of interference competition needs to be sufficiently high before the interactive selection of mass is stronger than the quality-quantity trade-off selection against mass.

The level of interactive competition that is required for this selection depends on an abundance that is so large that individuals meet sufficiently often in interactive competition, and this abundance depends on population growth with a quality-quantity balance that produces sufficiently many offspring from the net energy that is allocated to reproduction. The result is a population dynamic feedback attractor that selects mass in proportion to net energy by maintaining the level of interference competition that is needed to balance the selection of the quality-quantity trade-off (Witting 1997; with the selection attractor of invariant interference called a competitive interaction fixpoint).

This feedback selection indicates that the net energy that is used on replication could be a primary driver of natural selection. Net energy, however, is not a completely independent trait, because a product between resource handling and the pace of handling defines it (Witting 2017a,b; with population dynamic feedback selection selecting the pace of handling as the pace of metabolism, in proportion to mass-specific metabolism).

A secondary mass-rescaling selection that occurs during the feedback selection of mass is another essential factor to consider in relation to the hierarchy of natural selection causalities (Witting 2017a). A potential selection increase in mass implies that the larger offspring metabolises more energy during the period of parental care, and variants that avoid this extra metabolic cost of the extra mass will be selected over variants that do not. This selects variants that reduce the metabolic need by a decline in mass-specific metabolism, generating the observed allometric downscaling of mass-specific metabolism with mass.

The available net energy per unit physical time, however, declines with a decline in mass-specific metabolism reducing the reproductive rate. This aggregated problem of selecting mass with mass-rescaled metabolism is solved by variants that dilate biological periods and thereby maintains the net energy and reproduction of the organism on the per-generation timescale of natural selection. This generates the observed inverse allometric scaling between periods/ages and mass-specific metabolism.

The numerical response of this mass-rescaling selection is captured by the exponents of the body mass allometries. These are selected by the optimal foraging that generates the net energy for the overall feedback selection of mass, extending allometric scaling to ecological traits like home range and abundance. The result is a joint allometric scaling—of the metabolism, life history, and population ecology—that evolves as a sub-component of the natural selection of mass, instead of being a response to a physiological adaptation to size. For the original mathematical deductions of the allometric exponents see Witting (1995), for an extended deduction with primary selected mass-specific metabolism see Witting (2017a), and for a graphical deduction see Witting (2023).

Resource handling is one of the few life history traits that are unaffected by mass-rescaling selection (Witting 2017a). This makes it evolutionarily independent of mass, with a primary selection that drives the evolution of other traits by its contribution to the net energy driven population dynamic feedback selection.

Inter-specific variation in resource handling should thus explain large amounts of the variation in net energy and body mass, and secondarily also of the variation in other traits by their mass-rescaling dependence on the explained variation in mass.

Having removed the variance components that follow from variation in primary selected resource handling, I turn to the influence that the residual variation in the survival of offspring and adults have on the remaining life history variation. Ecological variation in mortality selects additional life history variation by perturbations of the competitive interaction fixpoint. An increase in mortality generates a decline in abundance and interactive competition, generating selection for increased replication until the interactive competition of the competitive interaction fixpoint is re-established. The energy for the selected increase in replication is taken primarily from body mass, with associated mass-rescaling selection for a wider range of life history variation.

Mass-specific metabolism is another potential life history influencer, as it is selected not only by secondary mass-rescaling but also by the primary selection that generates net energy for self-replication. The latter contributes to the feed-back selection of mass with superimposed mass-rescaling that downscales— at least to some degree—the primary selected mass-specific metabolism (Witting 2017a).

The importance of primary selected mass-specific metabolism for the selection of net energy and body mass is reflected in the values of the selected body mass allometries (Witting 2017a). Yet, birds have approximate Kleiber (1932) scaling with typical 1*/*4-like exponents, and this agrees with a theoretical prediction where it is the variation in resource handling (and not metabolic pace) that generates the variation in the naturally selected body masses. This seems to be the case for the majority of multicellular animals, while the allometric scaling of unicellular eucaryotes and especially prokaryotes indicates a major influence from primary selected metabolism in these taxa (Witting 2017a,b). Yet, to check for a potential influence from primary selected metabolism also in birds, following the decomposition from resource handling and ecological variation in mortality, I examine for a residual influence from mass-specific metabolism.

Having accounted for the selection influence from variation in net energy and mortality, I turn to the influence that the residual variation in the selection attractor of interactive competition has on the residual variation in other traits. With the most invariant component of the attractor being the intra-population fitness gradient in the cost of interactive competition, we expect some variation in our measure of interactive competition. Hence, the ecological traits that determine the level of interference (like abundance and home range overlap) should be selected to match the selected interference.

## 2 Methods

Following the selection hypothesis above, I decompose the inter-specific life history and ecological variation that Witting (2024) estimated for 11,187 species of birds, covering the parameters in Table 1. This variation was estimated from 37,305 published trait estimates, with the inter-specific extrapolations of missing parameters following from the allometric correlations of the data.

**Table 1:** Traits. The analysed traits, their symbols (S), units, and order of magnitude (M) inter-specific variation with medians, 5th, and 95th quantiles. Estimates from Witting (2024).

| Trait | S | Unit | M | 5th | Median | 95th |
| --- | --- | --- | --- | --- | --- | --- |
| Body mass | $w$ | kg | 4.8 | 0.007 | 0.036 | 1 |
| Res. handling | $\alpha$ | J | 7 | 0.00014 | 0.0016 | 0.16 |
| F. metabolism | $\beta$ | W/kg | 2 | 7.2 | 34 | 81 |
| B. metabolism | $\underline{\beta}$ | W/kg | 2.3 | 3.7 | 13 | 32 |
| Net energy | $\epsilon$ | W | 5.4 | 0.0089 | 0.058 | 1.4 |
| Incubation | $t_p$ | y | 1.3 | 0.032 | 0.041 | 0.087 |
| Fledging | $t_j$ | y | 2 | 0.029 | 0.047 | 0.16 |
| Rep. maturity | $t_m$ | y | 1.9 | 0.79 | 1 | 2.8 |
| Rep. period | $t_r$ | y | 1.7 | 1 | 1.8 | 4.2 |
| Generation | $t_g$ | y | 1.9 | 1.7 | 2.5 | 5.7 |
| Lifespan | $t_l$ | y | 1.7 | 6.1 | 10 | 26 |
| Off. survival | $l_m$ | $/t_m$ | 1.6 | 0.2 | 0.33 | 0.47 |
| Ad. mortality | $q_{ad}$ | $/y$ | 1.9 | 0.14 | 0.36 | 0.55 |
| Rep. rate | $m$ | $/y$ | 2 | 1.3 | 3.3 | 8 |
| Lifetime Rep. | $R$ | $/t_r$ | 1.6 | 4.3 | 6 | 9.9 |
| Abundance | $N$ | $/\text{km}^2$ | 5 | 1.3 | 14 | 130 |
| Biomass | $b$ | $\text{kg}/\text{km}^2$ | 5.2 | 0.051 | 0.73 | 7.9 |
| Pop. ene. use | $\epsilon_n$ | $\text{W}/\text{km}^2$ | 4 | 2.3 | 24 | 170 |
| Home range | $h$ | $\text{km}^2$ | 6.4 | 0.011 | 0.084 | 13 |
| Ho. overlap | $h_o$ | - | 6.5 | 0.14 | 1.8 | 110 |
| Interference | $\iota$ | - | 4.5 | -1 | 1 | 3.4 |

Some of the relevant traits are not available as data, and they were therefore calculated from other traits based on the trait relations in the population ecological model (Witting 2024). The variance decompositions that involve such traits are therefore not always independent of other traits, and this is discussed in the result section when relevant.

### 2.1 Explaining variance

I aim to explain the inter-specific variation within and across the 36 orders of birds. To calculate how variation in net energy (*α & β*), mortality (*q*_*ad*_ *& l*_*m*_), and interactive competition (*ι*) explain the inter-specific variation, I use double logarithmic relations 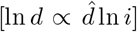 where exponents 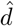 define the dependence of dependent traits *d* on the independent traits *i* ∈ *{α, q*_*ad*_, *l*_*m*_, *β, ι}*.

To predict a value 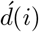, and calculate the associated residual value 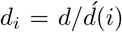, of a dependent trait *d* of a species in order *o*, I use a relation

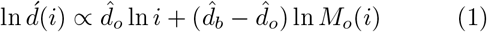

where *i* is the independent trait of the species, *M*_*o*_(*i*) is the median of *i* across the species in order *o*, 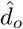 is the exponent that minimises the residual variance of ln 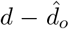 ln *i* across the species in the order, and 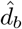 is the exponent that minimised the residual variance of ln 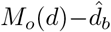 ln *M*_*o*_(*i*) across the medians of the different orders. The within order exponents 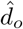 are estimated separately for all orders with life history estimates for more than *n* = 25 species, and set to the *n* weighted average of those within order exponents for orders with fewer estimates.

I present the values, and residual values, of traits as relative values 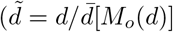 and 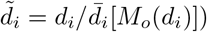 that are scaled by the average of the medians of the different orders, with residual values being calculated for the following sequence *α, q*_*ad*_, *l*_*m*_, *β*, and *ι* of the independent traits. The predictions are then evaluated by the average within-order variance of the dependent traits 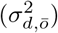 and their residuals 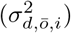, together with the between-order variance 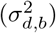, and residual variance 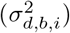, between the medians of the different orders.

To analyse the explained variance across traits, I use the proportion

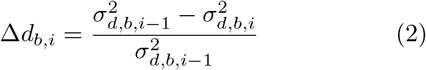

of the residual between-order variance in the dependent trait *d* that is explained by the independent trait *i*, and the corresponding proportion

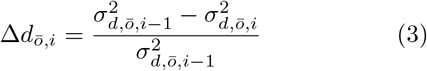

for the within-order variance. The total (*t*) proportion of the variance that is explained by all independent traits are 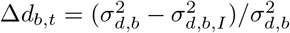 and 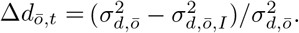

To analyse for differences in the variance that is explained between and within orders, I use the proportional difference

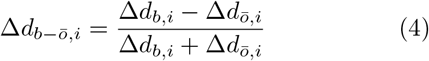

between the explained between and within order variance, with positive values implying that more variance is explained between orders than within, and negative values implying the opposite.

## 3 Results

Table 3 lists the estimated exponents (the average between the between order exponent and the *n* weighted average of the within order exponents) and the reduction in the within and across order variance of the different traits as a function of the independent trait components of *α, q*_*ad*_, *l*_*m*_, *β*, and *ι*. Fig. 1 illustrates these changes in the trait distributions of the different orders as the variance is explained by the independent traits (including for clarity only cases where an independent trait explains 10% or more of the variance of a trait). The widths of the distributions reduce, and the medians of the different orders converge on the overall median, as the independent traits explain the variance.

**Figure 1:**
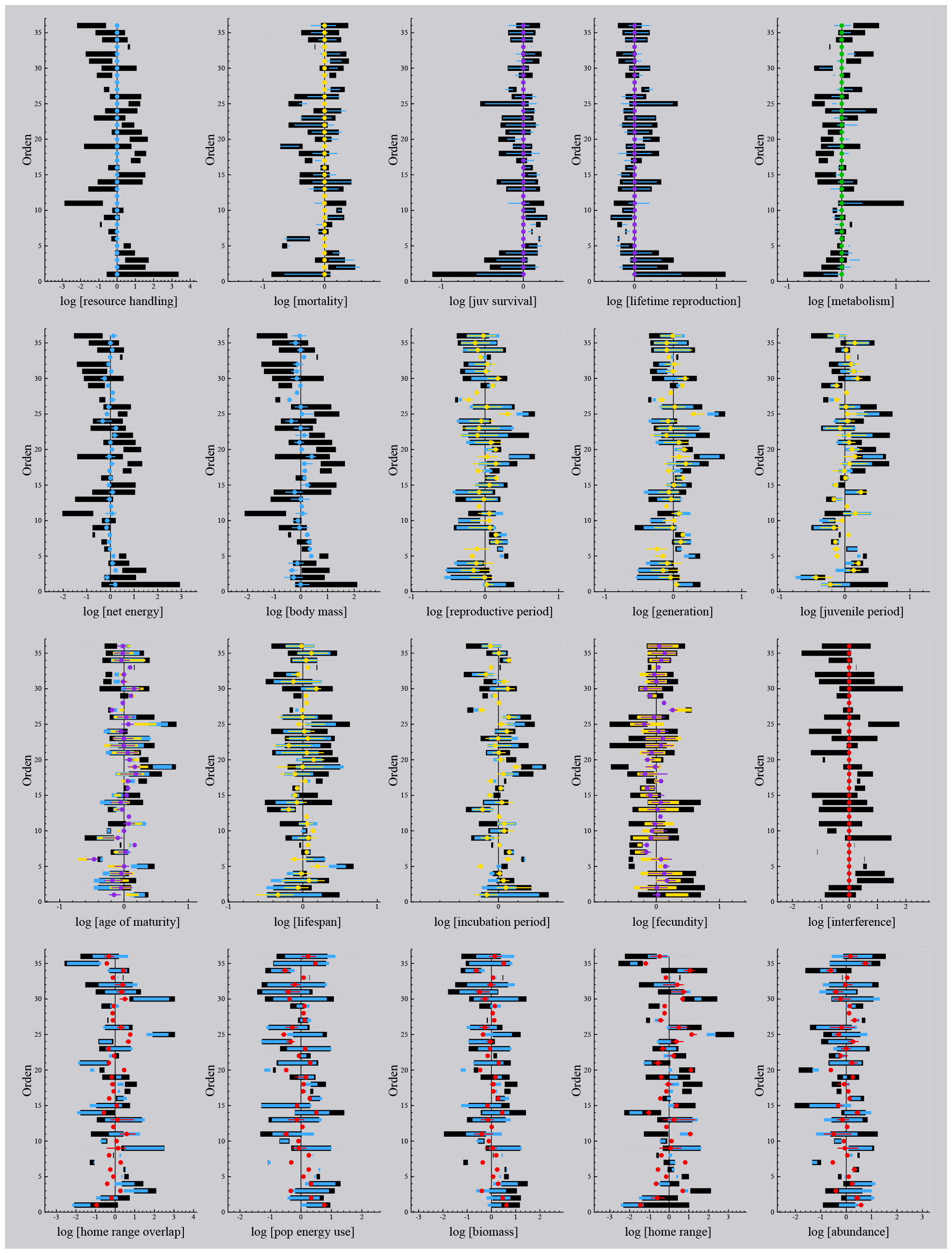
Explained variation. The 90% intervals (black bars), and limits (dashed lines), of the different traits across orders, including also the residual intervals and limits of the variation that is unexplained by resource handling (blue), residual adult mortality (yellow), residual juvenile survival (purple), residual metabolism (green), and residual interactive competition (red). Only cases where a dependent trait explains 10% or more of the variation are shown, with coloured dots being the medians of the residual variation for the last independent trait that explains more than 10%. Order: 1:Struthioniformes, 2:Galliformes, 3:Anseriformes, 4:Podicipediformes, 5:Phoenicopteriformes, 6:Phaethontiformes, 7:Eurypygiformes, 8:Mesitornithiformes, 9:Columbiformes, 10:Pterocliformes, 11:Caprimulgiformes, 12:Opisthocomiformes, 13:Cuculiformes, 14:Gruiformes, 15:Otidiformes, 16:Musophagiformes, 17:Gaviiformes, 18:Sphenisciformes, 19:Procellariiformes, 20:Ciconiiformes, 21:Pelecaniformes, 22:Suliformes, 23:Charadriiformes, 24:Strigiformes, 25:Cathartiformes, 26:Accipitriformes, 27:Coliiformes, 28:Leptosomiformes, 29:Trogoniformes, 30:Bucerotiformes, 31:Coraciiformes, 32:Piciformes, 33:Cariamiformes, 34:Falconiformes, 35:Psittaciformes, 36:Passeriformes

There is no overall tendency for my analysis to predict variation at one of the two taxonomic levels better than variation at the other (Table 2). The average amount of the deviations in the trait medians of an order (from the overall trait medians) that are explained by the independent traits is 72%, which is about similar to an average value of 76% for the within order explained variance.

**Table 2:**
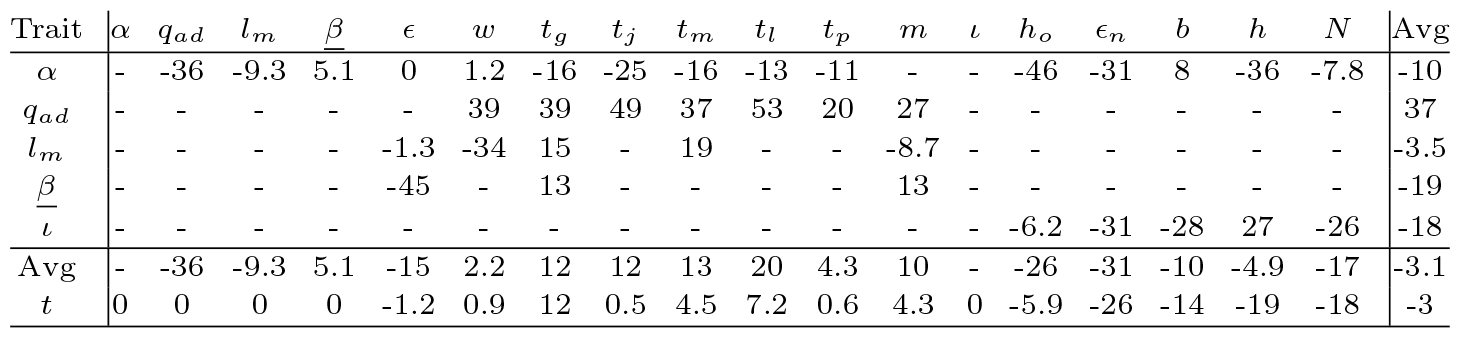
Between versus within order variance. The proportional difference (eqn 4, in percent) between the between and within order variance that is explained by the independent traits. Positive values are cases where more between order variance than within order variance is explained, and negative values are cases where more within than between order variance is explained. Calculated only for dependent traits where more than ten percent of the total variance is explained by an independent trait, with *t* denoting the joint effect of all independent traits *i* ∈ *{α, q*_*ad*_, *l*_*m*_, *β, ι}*.

**Table 3:**
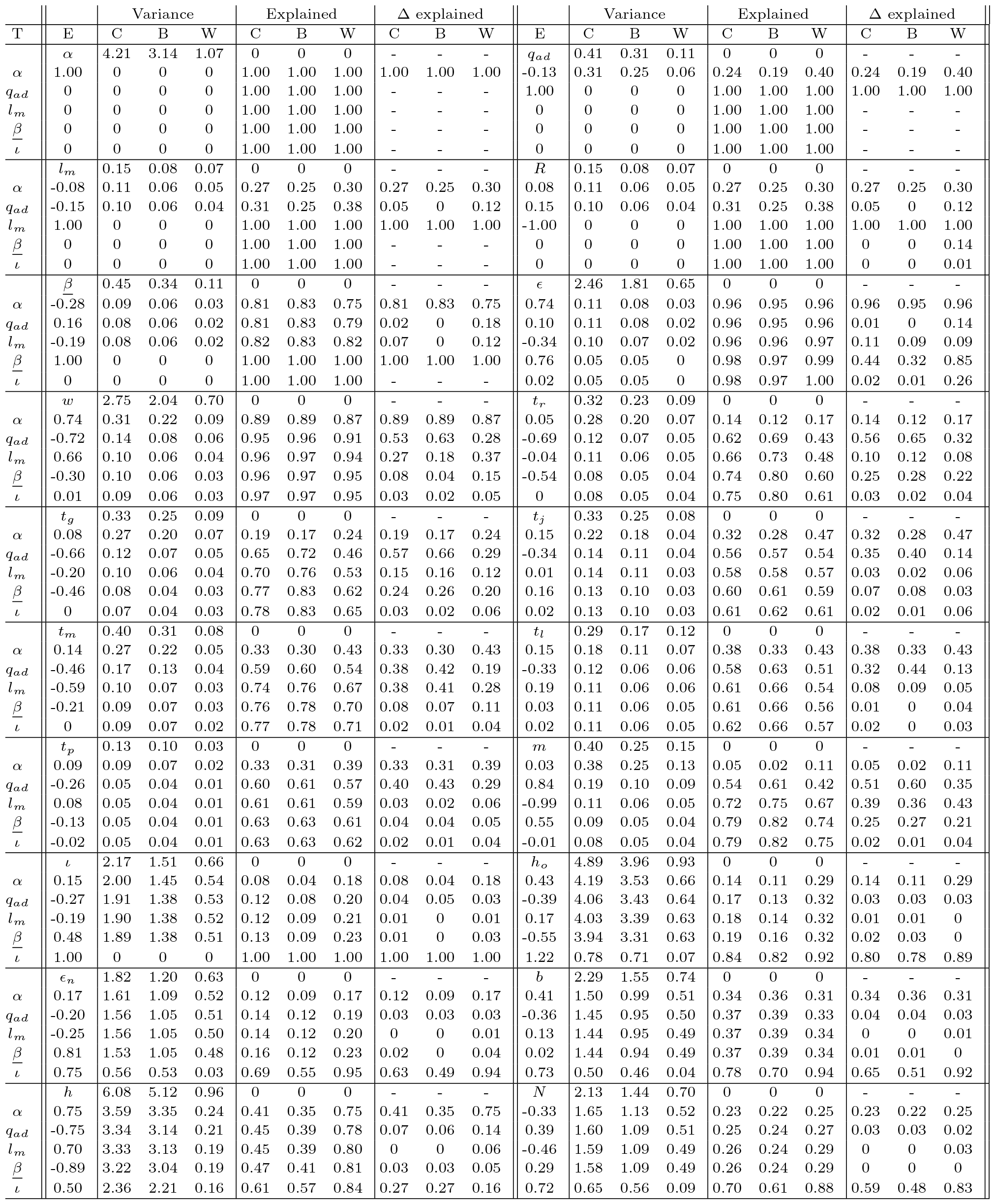
Explained variance. Variance: First rows: The between orden (B) variance, average within orden (W) variance, and combined variance (C=B+W) for the trait in columns E. Other rows: The residual variance that is not explained by allometric correlations with the traits in column T. Explained: The fraction of the total variance that is explained. Δ explained: The fraction of the residual variance that is explained by the column T trait (eqns 2 and 3). The E column values are the average of the between and within order exponents that explain most variance.

### 3.1 Selection decomposition

#### Resource handling

Resource handling (*α*) generates net energy (*c*) for the population dynamic feedback selection of mass, with the associated mass-rescaling selection inducing secondary effects on other traits (Witting 2017a). My estimates of resource handling, however, are estimated from net energy and mass-specific metabolism (*α* = *c/β*), and net energy is estimated from the combustion energy of body mass (*w*_*e*_), yearly reproduction (*m*), and metabolism 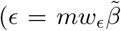, with 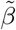 being a relative measure of the energy that is metabolised by offspring; Witting 2024). The variance decomposition from resource handling should thus not be seen as a statistical test, but only as a description of how the population ecological model reconciles the inter-specific life history variation with the different components of the net energy that generates the selection of the life histories.

For birds, where the empirical allometries fit with body mass variation that is selected predominantly from variation in resource handling (Witting 2017a), net energy (*c*) is predicted to scale as a ln *c* ∝ 0.75 ln *α* function of resource handling, with body mass selected in proportion with resource handling (ln *w* ∝ 1.33 ln *c* ∝ 1 ln *α*), and the second component of net energy 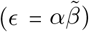, i.e., metabolic pace 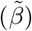, being a declining function of resource handling 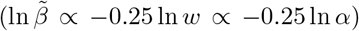 owing to the selected mass-rescaling decline in mass-specific metabolism (ln *β* ∝ −0.25 ln *w*).

This is reflected in the variance decomposition of the life history models, where resource handling raised to the 0.74 power accounts for 96% of the variance in net energy. 89% of the variance in body mass follows from a somewhat lower than proportional dependence on resource handling (0.74 exponent), and 81% of the variance in mass-specific metabolism is captured by resource handling raised to the −0.28 power.

Somewhat lower percentages between 32% and 41% are explained for four periods/ages (*t*_*p*_, *t*_*j*_, *t*_*m*_, *t*_*l*_), plus biomass and home range, and about 20% of the variance is explained for adult mortality, offspring survival, lifetime reproduction, generation time, and abundance. It is only for the reproductive rate and level of interference, that less than 10% of the variance is explained by resource handling.

These relations tend to follow the expected secondary effects of mass-rescaling. Given the 0.74 power dependence of body mass on resource handling, the expected and observed exponents are −0.25 * 0.74 = −0.18 and −0.13 for adult mortality, 1 * 0.74 = 0.74 and 0.75 for the home range area, −0.75 * 0.74 = −0.55 and −0.33 for the abundance of populations, and 0.25*0.74 = 0.18 and about 0.15 for three of the best explained life history ages/periods. For the expected invariance of lifetime reproduction and the probability that an offspring survives to the reproductive age, we find a rather week dependence with average exponents of 0.08 and −0.08. Yearly reproduction is somewhat surprisingly almost invariant of resource handling with an observed exponent of 0.03.

#### Adult mortality

Variation in mortality affects the selection of several traits by a perturbation of the selection attractor of interactive competition (Witting 1997, 2008). Increased mortality selects for an increase in reproduction that compensates for the decline in abundance and interference competition that follows from increased mortality. This selection affects several other traits secondarily, as the net energy for increased reproduction is generated predominately from a selection decline in mass; with the associated mass-rescaling having potential effects on other traits.

This selection is reflected in the residual variation that is explained by the residual variation in adult mortality. 51% of the residual variation in the annual rate of reproduction, and 53% of the residual variation in mass, are explained by the residual variation in annual mortality, with the rate of reproduction increasing (estimated exponent of 0.84) and the body mass declining (estimated exponent of -0.72) with increased mortality. A cascading mass-rescaling effect—with shorter life periods from the smaller masses of increased morality— was also found. The residual variation in adult mortality explains 40% (-0.26 exponent) of the residual variation in the incubation period, 35% (-0.34 exponent) of the juvenile period, 38% (-0.46 exponent) of the age of maturity, and 32% (-0.33 exponent) of the variation in lifespan (here you should not pay attention to *t*_*r*_ and *t*_*g*_ as these are partially calculated from *q*_*ad*_). No massrescaling effect, however, was found for mass-specific metabolism.

With the explained levels of the residual variation being below seven percent, the ecological traits of abundance, home range overlap, and level of interference are largely unaffected by the residual variation in mortality. This may come as a surprise, as increased mortality is usually assumed to have a direct negative impact on the abundance of populations. Yet, the latter response is a population dynamic response in the absence of evolutionary changes, while the selection dynamics of the competitive interaction fixpoint compensate for the decline in abundance and interference, by the selection of net energy from mass to reproduction until the abundance and level of interactive competition of the evolutionary equilibrium is re-established.

The expected changes in abundance and home range to changes in mortality are the secondary mass-rescaling effects from the associated evolutionary changes in mass. The abundance is thus predicted to increase, and the home range to decline, with a mass that is selected to decline from increased mortality. This second order response, however, was not really detected with only 7% and 3% of the residual variance in home range and abundance being explained by the residual variation in adult mortality; yet the directions of the responses were as expected with a negative (−0.75) exponent for home range and a positive (0.39) for the abundance of populations.

#### Offspring survival

The probability (*l*_*m*_) that an offspring will survive to the age of reproductive maturity is predicted to have a large impact on fecundity (*m*) and lifetime reproduction (*R* = *t*_*r*_*m*) through the survival versus reproduction compensation in population dynamic feedback selection. Yet, the potential effect *R* = 2*/l*_*m*_ is incorporated as a constraint in the estimated models given the assumption of stable populations, where *l*_*m*_*R* = 2 is the expected lifetime reproduction of a female.

The feedback selection compensation between survival and reproduction, however, operates through a reallocation of energy between reproduction and mass. As for adult survival, we find that increased offspring survival selects for an increase in body mass, with an estimated exponent of 0.66, and 27% of the residual variation in body mass explained by the residual variation in offspring survival.

With less than 10% of the residual variation explained for most of the remaining traits, a secondary mass-rescaling effect is not really detected. There is instead a negative relation (-0.59 exponent) between *l*_*m*_ and reproductive maturity, with 38% of the residual variation is explained, most likely, from a probability of survival that declines the longer offspring need to survive to reach the reproductive age.

#### Metabolism

Mass-specific metabolism affects rate-dependent traits, including the pace of resource handling that generates net energy for the selection of mass. With 82% of the variation in mass-specific metabolism being explained already by mass-rescaling from primary variation in net energy and mortality, there is some residual metabolic variation left to detect extra variation in net energy and mass. Yet, with 96% of the variation in net energy explained already by resource handling, we can expect only a very small influence on the life history from primary (i.e., non mass-rescaling selected) variation in mass-specific metabolism.

This is reflected in the variance decomposition, where mass-specific metabolism explains 44% of the residual variation in net energy, with the dependence being about proportional as expected (exponent of 0.76). Yet, while the dependence of residual variation in net energy on mass-specific metabolism is strong, primary selection on mass-specific metabolism accounts for no more than 2% of the total variation in net energy. The small but positive increase in net energy is not detected in body mass, where the exponent is negative (−0.30) and only 8% of the residual variation is explained.

Nevertheless, several rate-dependent traits have about 25% of their residual variation explained by mass-specific metabolism, with the direction of the response being as expected, i.e., rates correlate positively [exponent of 0.55 for annual reproduction] and periods and ages negatively [exponents of −0.54 for reproductive period, −0.46 for generation time, and −0.21 for reproductive maturity] with mass-specific metabolism.

#### Interactive competition

The level of intra-specific interference per individual (*ι*) is probably the most derived of all the traits considered, in the sense that the mechanistic generation of interference depends on a multitude of ecological and physiological traits that change with the evolutionary modification of the life history (approximated here by calculating interference as a function of abundance, home range, metabolism, and body mass). But, instead of being a derived trait that follows as a passive consequence of natural selection changes in other traits, the level of interference is one of the most central independent traits in population dynamic feedback selection. It is the overall selection attractor that controls the natural selection of the life history by selecting net assimilated energy between the demographic traits and mass, with the attractor itself being unaffected by the selected variation.

The attractor is referred to as the competitive interaction fixpoint, and it has a theoretical interference level of *ι*^**^ = 1*/ψ* when body mass in selected at an evolutionary equilibrium, and a theoretical level of *ι*^\**s*^ = (4*d* − 1)*/ψ*(2*d* − 1) when mass is selected to increase exponentially at an evolutionary steady state [Witting, 1997; *ψ* is the intra-population gradient (around the average life history) in the cost of interference (e.g., different access to resources) per unit interference on log scale; subscripts ** and \**s* denote evolutionary equilibrium and steady state; *d* is the dominant number of spatial dimensions in the foraging ecology of birds]. The real selection invariant parameter is not the level of interference itself, but the intrapopulation gradient in the cost of interference [*ι*^**^*ψ* = 1 & *ι*^\**s*^*ψ* = (4*d*−1)*/*(2*d*−1)]. This gradient—that favours the large and competitively superior individuals in the population—is selected to balance the fitness gradient of the quality-quantity trade-off.

With unconstrained selection being more likely to occur among the larger and ecologically dominant species, there might be some positive correlation where a larger fraction of the species that are situated at the *ι*^\**s*^ attractor are those with the largest resource handling and/or metabolic pace. Yet, apart from this potential correlation, we expect a level of interference that is invariant with respect to the selected variation in the life history. This is also to a large degree what we find.

Even though the level of interference is calculated from abundance, home range, metabolism, and body mass, it is the most invariant traits, i.e., the trait that is least affected by the variation in net energy and mortality. Where 26, 47, 96, and 82% of the total variance in abundance, home range, body mass and metabolism have been explained so far by independent traits, only 13% of interactive competition is explained. It is evident that a large fraction of the explained variation in the sub-components cancel out, leaving the level of interference largely unaffected by the underlying variation in other traits.

Resource handling and mass-specific metabolism account for 8% and 1% of the variation in interactive competition, with a positive dependence (exponents of 0.15 and 0.48) as expected from the likely link between net energy and the likelihood of unconstrained selection. Four extra percent is reconciled by adult mortality, with a negative dependence (exponent of -0.27). While the level of interference should in principle be independent of mortality, the direction of the response (where the level of interference is declining with an increase in mortality) indicates that it may take some time for the selection attractor to adjust to ecological changes in the rate of mortality.

As expected from the invariance, the residual variation in the estimated level of interference explains three percent or less of the residual variation in all of the demographic and physiological traits, including mass. Yet, with the variation in population density and home range being selected to match the level of interference at the selection attractor, we find that the residual variation in interactive competition reconciles 27% of the residual variance in the home range, 59% for population density, 63% for the energy use of populations, 65% for biomass, and 80% for the overlap between home ranges. Of these ecological factors, it is especially the home range overlap that relates most directly with the level of interference, as the probability to encounter other individuals is a direct function of the degree of overlap between home ranges.

## 4 Discussion

Body mass is an essential evolutionary player that influences the evolution of other traits, but it is not the primary driver of natural selection as its selection depends on other traits. I found variation in resource handling and mortality to reconcile 96% of the body mass variation in birds. The associated secondary mass-rescaling explained 74% of the variation in metabolism, reproduction, and life periods/ages, and 36% of the variation in abundance and home range. All life history traits including body mass and metabolism, had no more than 1% of their variation explained by interactive competition, which explained 80% and 59% of the residual variation in home range overlap and abundance. No consistent mass and mass-rescaling response was detected from the residual variation in mass-specific metabolism, confirming that it is primarily variation in resource handling that generates the body mass variation and allometric scaling of birds.

The observed negative dependence of body mass on adult (*q*_*ad*_ exponent of -0.72) and offspring (1−*l*_*m*_ exponent of −0.66) mortality is documented in other studies, mainly for fish (Reznick et al. 1996; Haugen and Vøllestad 2001; Sinclair et al. 2002; Carlson et al. 2007; Herczeg et al. 2009). It supports a central mechanism, where population dynamic feedback selection allocates net energy between reproduction and mass to maintain the interference of the competitive interaction fixpoint. One implication of this is the absence of a decline in abundance following increased mortality. The predicted, and weakly observed, correlation is instead an increase in abundance with increased mortality, reflecting the secondary mass-rescaling that follows from the selected decline in mass following increased mortality. In conclusion, I found population dynamic feedback selection to be a better predictor than density regulation for the observed inter-specific covariance between mortality and abundance.

Being based on the bottom-up unfolding of natural selection, my study is not comparable with comparative analyses that use life history correlations with body mass, mortality, lifestyles, energy use, and phylogeny to generate hypotheses of life history evolution (e.g. Promilsow and Harvey 1990; Blackburn 1991; Sæther and Bakke 2000; Bielby et al. 2007; Dobson and Oli 2007; Brown et al. 2018; Burger et al. 2019). While these studies identify inter-specific trait correlations that follow from evolution, they do not necessarily identify the underlying natural selection causes. Comparative approaches tend to assume that variation in mass is the primary cause for evolutionary variation in other traits. Yet, by analysing the causality of natural selection I found the selection of body mass to be a secondary effect from the primary selection of resource handling and metabolism, combined with ecological variation in resource density and mortality.

The reopening (Dobson 2012) of historical/phylogenetic ecology (e.g. Holder 1983; Brooks and McLennan 1991; McKitrick 1993; Brown 1994) by Sibly et al. (2012) as a proposed essential component for bird life history evolution is also not a real search for cause and effect in natural selection. Being the fundament of pre-Darwinian classification (Linnaeus 1758), the observation that the diversity of life is better accounted for by grouping organisms into more closely related taxa is ancient. While phylogeny reflects evolutionary diversification under the Darwinian hypothesis, and while it identifies life history correlations among related species, historical ecology and comparative phylogeny provides no essential insights into the underlying natural selection causes behind the evolved life history variation (Reeve and Sherman 2001). Life history differences by phylogenetic distance is the evolutionary outcome of natural selection and other processes of evolution, and not the cause of evolution.

Where comparative methods and historical ecology represent non-causal correlation analyses, Lack’s (1947) clutch size is one of the first causal natural selection analyses. By quantifying observed trade-offs between the current reproductive effort of parents and the future survival of parents and offspring, the original and subsequent analyses documented widespread selection towards intermediate clutch sizes (Lack 1947; Charnov and Krebs 1974; Schaffer 1983; Boyce and Perrins 1987; Daan et al. 1990; Godfray et al. 1991; Ylönen et al. 1998).

Lack’s clutch size appears to predict the reproductive rate from the trade-off between the current reproductive effort and future survival. But the mathematical selection models behind Lack’s clutch size do not document the bottom-up natural selection stability of the trade-off. When the applied framework of constant relative fitnesses is taken literally, the predicted clutch size is evolutionary unstable because the essential tradeoff is scale-dependent in such a way that the quality-quantity trade-off selects for a decline in the amount of energy that is allocated by the trade-off (Witting 1997, 2008). This reflects a deeper selection for a continued increase in reproduction by a corresponding decline in mass.

To stabilise this selection, we need a frequencydependent selection of mass to balance the frequency-independent quality-quantity selection against mass. The population dynamic feedback of density dependent interactive competition provides this balance, selecting an overall balance between reproduction and mass from the net energy and individual mortality of the species. Superimposed upon this there is a more derived selection that optimises the physiology, and this is expected to continue until the trade-off between the current reproductive effort and future survival matches the net energy, mass, and reproductive rate of the competitive interaction fixpoint (Witting 1997, 2008). Hence, the top-down backwards selection analysis of Lack’s clutch size meets the bottom-up forward selection of Malthusian relativity, providing a more unified theory where the traditionally assumed contingency follows from a natural selection that unfolds from the origin of replicating molecules.

